# Proteome-wide structure-based accessibility analysis of ligandable and detectable cysteines in chemoproteomic datasets

**DOI:** 10.1101/2022.12.12.518491

**Authors:** Matthew E. H. White, Jesús Gil, Edward W. Tate

## Abstract

Covalent drug discovery, in particular targeting reactive cysteines, has undergone a resurgence over the past two decades, demonstrated by recent clinical successes of covalent inhibitors for high-priority cancer targets. Reactive cysteine profiling, first pioneered by the Cravatt lab, has emerged in parallel as a powerful approach for proteome-wide on- and off-target profiling. Thus far however, structural analysis of liganded cysteines has been restricted to experimentally determined protein structures. We combined AlphaFold-predicted amino acid side chain accessibilities for >95% of the human proteome with a meta-analysis of thirteen public cysteine profiling datasets, totalling 40,070 unique cysteine residues, revealing accessibility biases in sampled cysteines primarily dictated by warhead chemistry. Analysis of >3.5 million cysteine-fragment interactions further suggests that exposed cysteine residues are preferentially targeted by elaborated fragments and drug-like compounds. We finally propose a framework for benchmarking coverage of ligandable cysteines in future cysteine profiling approaches, considering both selectivity for high-priority residues and quantitative depth. All analysis and produced resources (freely available at www.github.com/TateLab) are readily extendable to reactive amino acids beyond cysteine, and related questions in chemical biology.

## Introduction

Covalent drug discovery has re-emerged over the past two decades as a powerful modality for difficult-to-drug and conventionally ‘intractable’ targets. Covalent inhibition of protein targets takes advantage of the inherent reactivity of specific amino acid side-chains, primarily cysteine, but with a continually expanding scope encompassing lysine, threonine, histidine and even electrophilic N-terminal modifications.^1–5^ Covalent binding by a targeted covalent inhibitor (TCI) presents a number of advantages over non-covalent binding, including extended rather than equilibrium-limited residence time at a target site, potentially increased tractability of shallow binding pockets or intrinsically disordered regions and selectivity based on disease-associated amino acid mutations (Figure 1a).^6,7^ These benefits are exemplified by development and FDA-approval of covalent inhibitors for high-priority cancer targets, including KRAS[G12C]^7,8^, EGFR^9^ and BTK^10^. Even covalent ligands for amino acids outside of enzyme active sites may offer valuable starting points as allosteric inhibitors or for covalent bifunctional molecules which recruit effector proteins to neosubstrates (e.g. covalent PROTACs^11,12^ or DUBTACs^13^). However, systematic discovery of novel and developable covalent ligands remains a significant bottleneck due to the requirement for balanced reactivity and sufficient selectivity towards a single target amino acid against all other accessible amino acids displaying similar chemistry.

**Figure 1:**
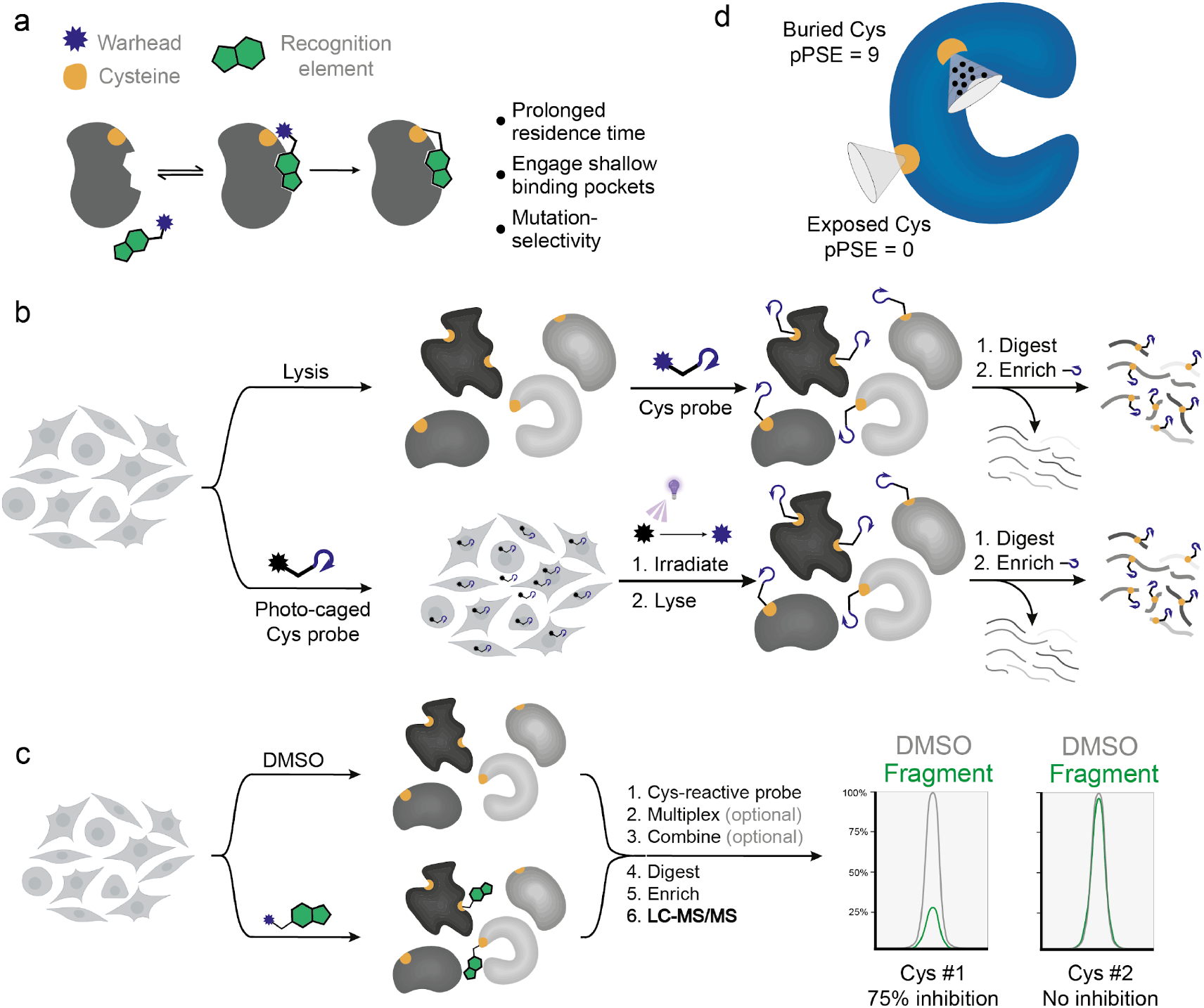
Overview of cysteine profiling approaches and AlphaFold2-based accessibility analysis applied in this study. **A.** Representative mode-of-action for covalent binding by a targeted covalent inhibitor. **B.** Schematic showing cysteine profiling approaches applying peptide-level cysteine enrichment, either by labelling in lysates (top) or in live cells (bottom). **c.** Overview of competitive fragment screening by mass spectrometry-based cysteine profiling, with representative chromatograms of fragment-competed and non-competed cysteines shown. **d.** Accessibility analysis on AlphaFold-predicted structures as described by Bludau et al.^21^

Technology platforms which permit quantitative profiling of covalent protein modifications have become powerful tools for TCI discovery and development.^6^ Competition-based chemoproteomic methods such as <u>iso</u>topic <u>T</u>andem <u>O</u>rthogonal <u>P</u>roteolysis—<u>A</u>ctivity-<u>B</u>ased <u>P</u>rotein <u>P</u>rofiling (isoTOP-ABPP), initially developed by Cravatt and co-workers,^1^ have seen widespread application for highly parallel and versatile analysis of potential amino acid reactivity and ligandability. Such analyses initially focused on cysteine-reactive TCIs in cell lysates (*in vitro*) using a cysteine-reactive iodoacetamide warhead to enrich, identify and quantify peptides bearing a cysteine existing at least partially in a reactive, and therefore potentially druggable, state (Figure 1b, top).^1,2^ Subsequent technical developments, including novel cleavable reagents, multiplexing strategies, chemical enrichment and mass spectrometric (MS) acquisition approaches, have together markedly improved both depth and throughput, as reviewed comprehensively elsewhere.^14,15^ Weerapana and co-workers further developed this concept for live-cell (*in situ*) labelling, using photo-uncaging of protected α-haloketones in live cells to permit labelling concentrations of up to 200 μM without significant cell death (Figure 1b, bottom).^16,17^ Such reactive amino acid profiling platforms have been applied extensively to covalent ligand discovery, primarily through quantifying cysteine reactivity in lysates treated with libraries of electrophilic fragments compared to DMSO-treated controls (Figure 1c). In these screens, loss of signal for a cysteine-containing peptide in a fragment-treated sample is interpreted as evidence for covalent fragment reactivity at that residue, with the magnitude of signal loss proportional to the occupancy of this interaction. In recent years, increasingly large-scale fragment library screens have generated a substantial body of publicly available datasets.^2,15,18–20^

Whilst the most comprehensive cysteine profiling (CP) studies consistently profile 10,000 or more reactive cysteines in parallel, the structural distribution of these residues across the 261,260 cysteine residues (from the UniProt one-protein-per-gene database) present in the human proteome has yet to be systematically analysed. Similarly, the extent to which CP approaches sample potentially ligand-accessible residues remains undefined but has significant implications for the efficiency of TCI discovery platforms and assessment of proteome-wide selectivity. Here, we present a meta-analysis of 13 published reactive cysteine datasets (Table S1) in the context of proteome-wide residue accessibility predictions enabled by AlphaFold (Figure 1d). We find significant variation in the accessibility distributions of profiled cysteines across published studies, uncovering warhead-specific effects and clear disparities in residue targeting between cysteine-reactive enrichment probes, electrophilic fragment libraries and even optimised TCIs. These results will help inform the applicability of different CP approaches to various stages of the drug discovery pipeline, including direct screening by MS-based ABPP, target identification and off-target profiling, and suggest directions for future development of CP platforms to enhance and accelerate discovery of developable covalent ligands.

### Amino acid side chains and post-translational modifications have distinct accessibility profiles

To predict the accessibility of amino acid side chains we used an approach recently reported by Bludau et al. to integrate post-translational modification (PTM) proteomic datasets with AlphaFold-predicted protein structural information across almost all humans proteins.^21,22^ In their study, solvent accessibility calculations and predictions of folded/intrinsically-disordered regions were integrated with phosphorylation, ubiquitination and O-glycosylation datasets, and applied to uncover a number of PTM-specific structural distribution patterns. Of particular interest to the present study, the incorporation of AlphaFold prediction error into part-sphere exposure^23,24^ calculations from predicted structures provides a proteome-wide metric of side-chain solvent accessibility at single residue resolution. Applying this approach, we calculated ‘prediction-aware part-sphere exposure’ (pPSE) for each residue in 20042 proteins from the human UniProt sequence database. The pPSE value of a given amino acid reflects the number of proximal α-carbons counted in a conical volume projecting along the Cα-Cβ vector (or pseudo-vector in the case of glycine). A high pPSE value represents a crowded environment around the side chain and therefore a more buried and less accessible residue, whereas a low pPSE value indicates high accessibility.

We assessed the overall validity of this model across a range of relevant benchmarks. First, considering patterns of accessibility across individual amino acids, the intrinsic physicochemical and structural properties of certain amino acids show intuitive trends. For example, charged amino acids such as Asp, Glu, Lys and Arg show enrichment for highly exposed residues, whereas neutral and hydrophobic residues such as Val, Leu, Tyr and Phe show a converse enrichment for buried residues (Figure 2a). Both Gly and Pro show a very strong enrichment for fully exposed (pPSE = 0) but not partially exposed residues (1 < pPSE < 5), potentially reflective of their known enrichment in short loop regions.^25^ Whilst general trends in hydrophobicity can be extracted from proteome-wide averages, focusing on subsets of annotated cysteine PTMs also shows distinct distributions. For example, cysteines known to form disulfide bonds (by UniProt annotation^26^) show a broad distribution across all pPSE values, reflective of the structural nature of internal disulfides as well as prevalence of more exposed, redox-active cysteines. Conversely, lipid-modified cysteine residues showed a significant bias for highly exposed residues, with 94% of 481 *S*-acylated cysteines found at pPSE < 5 and 100% of 220 known prenylated cysteines (both from UniProt annotation^26^) at pPSE < 3 (Figure 2b). pPSE distributions with high exposure are consistent with the requirement for enzymatic lipidation at cysteine, and mediation of subsequent membrane interactions at the protein surface.^27^ Active site cysteine-annotated residues^26^ showed greatest enrichment at 5 < pPSE < 8, indicating the predicted ‘depth’ of annotated active sites strikes the expected balance between substrate accessibility and solvent exclusion. Together, these analyses indicate that physicochemically and biologically relevant trends for amino acid solvent accessibility can be extracted from computationally predicted structures alongside conservative filtering for prediction quality. However, we note that these predictions remain subject to the limitations of AlphaFold itself, and caution should be exercised when drawing conclusions for any single residue in isolation in the absence of experimental validation.

**Figure 2:**
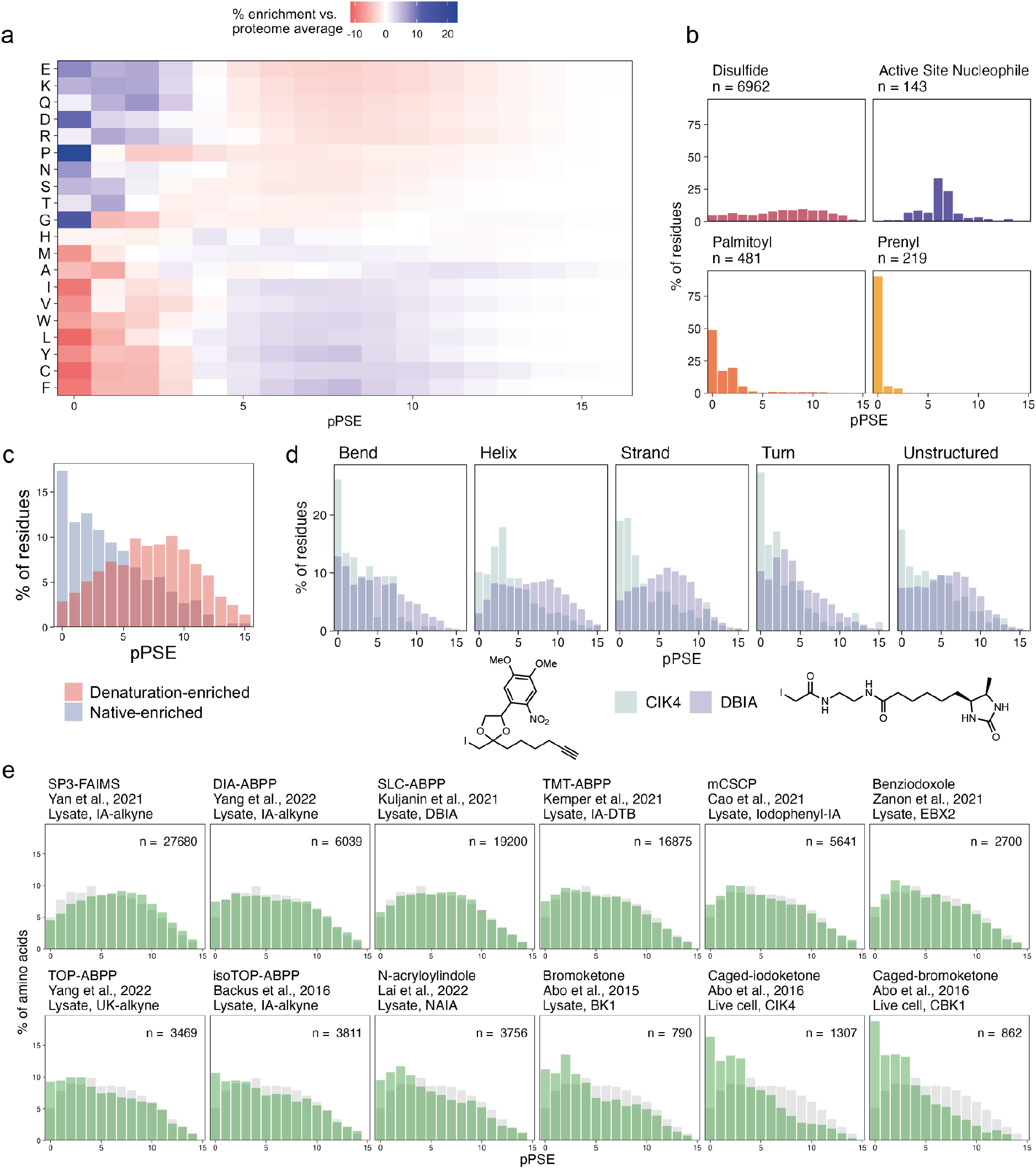
Cysteine profiling workflows sample significantly different predicted accessibility distributions by warhead chemistry and departure from native cellular conditions. **a.** Proteome-wide amino acid pPSE distributions - normalised to the whole-proteome average for all residues - reflect known physicochemical properties of side chains. Amino acids are ordered by percentage exposed (pPSE < 5) residues. **b.** pPSE distributions of cysteine residues with annotated PTMs, as described in UniProt. **c.** Effect of heat-denaturation on accessibility of profiled cysteines. Native-enriched residues (R_Native/Heat_ > 2, blue) and denaturation-enriched residues (R_Native/Heat_ < 0.5, red) show distinct accessibility distributions. Data from Backus et al.^2^ **d.** Comparison of iodoacetamide- and caged-bromoketone warhead across annotated secondary structure groups from Bludau et al.^21^ Data from Kuljanin et al.^18^ (desthiobiotin-iodoacetamide, DBIA) and Abo et al.^17^ (caged-iodoketone, CIK4). **e.** Accessibility distributions for 13 literature CP datasets (green) compared to the whole-proteome average distributions (light grey). Reference details, cysteine-reactive probe and labelling conditions are denoted in figure labels. Structures for each probe are shown below the plot.

### Warhead chemistry and labelling environment generate distinct accessibility profiles

The question of cysteine coverage remains important for CP workflows, as current acquisition approaches are limited to sampling a maximum of ~20,000 unique cysteines in a single run.^28^ Although this coverage represents a tremendous improvement over first-generation technology, it encompasses only 10% of all MS-detectable tryptic peptides and just 7.5% of all cysteines in the proteome.^28^ Clearly, the subset of therapeutically-relevant cysteines is significantly smaller, as a residue must be accessible to a small molecule, present at least partially in the nucleophilic thiol/thiolate form, and in the case of TCIs should result in a phenotypic change upon drug binding. As such, enrichment of the most ‘ligandable’ cysteines would achieve meaningful coverage of those residues most likely to prove fruitful as therapeutic targets, although there is currently no clear consensus on the size or character of this subset.

In isoTOP-ABPP, preferential enrichment for surface residues has been proposed to occur by labelling with activity-based probes in minimally denatured lysates. For example, Backus *et al*. performed a direct comparison of cysteines quantified in a ‘native lysate’ compared to those denatured by heating. Comparison of pPSE distributions between these two strongly contrasted conditions demonstrates that residues with at least two-fold higher enrichment in heat-treated samples (R_Native/Heat_ < 0.5) also show increased pPSE, suggesting increased access to more buried cysteines (Figure 2c). Conversely, cysteines at least two-fold enriched in the non-denatured lysate (R_Native/Heat_ > 2) are enriched in exposed (pPSE < 6) cysteines, confirming that minimally-denaturing lysis preserves exposed cysteine labelling preferentially and that heat-denaturation increases the accessibility of previously buried residues.

We next sought to understand how the subset of cysteines sampled by a range of published CP protocols varied by accessibility and secondary structure. We compiled 13 published datasets, profiling a total of 40,070 unique cysteine residues and >3,500,000 fragment-cysteine interactions across a range of warhead chemistries, enrichment strategies and MS acquisition approaches (Table S1). Among iodoacetamide-based reagents, the overall subset of cysteine residues sampled is remarkably similar to the background distribution of cysteines (Figure 2e). Notably, both N-acryloylindole (NAIA) and α-bromoketone (BK1) warheads showed modest enrichment of more exposed residues (Figure 2e, bottom middle panels). The clearest enrichment is observed with photocaged α-haloketones (Figure 2e, lower right panels), which permeate live cells without significant labelling and are then uncaged by prompt irradiation at 365 nm (as shown in Figure 1b, lower workflow).^16^ Although with markedly lower overall cysteine coverage (1497 unique cysteines in total across both live-cell datasets), live-cell labelling with photocaged warheads shows significant bias for exposed residues across all annotated protein secondary structures (Figure 2d), and thus does not arise merely due to labelling highly accessible, unstructured regions. The additional enrichment of accessible cysteines resulting from uncaging and live-cell labelling (over BK1 labelling in lysates) may be attributable to the native labelling cellular environment, as it is reasonable to suppose that even mild, detergent-free lysis may present a measurable departure from native cellular conditions, for example by oxidation of highly reactive residues^29^ or partial denaturation by sonication. Furthermore, the timescales of uncaging and labelling are potentially different in live cells compared to lysates, especially as probe quenching may be significantly higher in redox-buffered live cells. Finally, high concentrations of the labelling reagent may induce some level of denaturation upon multiple labelling events on a given protein. Taken together, these data indicate that probe chemistry and labelling conditions significantly affect the subset of cysteines sampled in a given experiment and should be critically evaluated based on the experimental design.

### Reactive fragments and drug-like covalent inhibitors primarily target accessible residues

Beyond cysteine coverage by enrichment, the size of the ligandable cysteinome is also largely undetermined. Although a number of computational studies have defined ligandable cysteines from analysis of cysteine orientation, adjacent residues and binding pocket characterisation^30–32^, these approaches have thus far relied on experimental structures. We therefore combined multiple fragment screening datasets with pPSE analysis to provide a proteome-wide window on potentially druggable cysteines. In the first report of covalent fragment screening by isoTOP-ABPP, a comparison of fragment treatments in lysates (*in vitro*) or live cells (*in situ*) was performed, showing the liganded cysteines in both treatments prioritise more accessible residue distributions when compared to all detected cysteines (Figure 3a). There also appears to be a greater bias in live cells treated with fragments, suggesting that these highly accessible cysteines represent the most ligandable covalent targets in a native cellular environment. Although significantly more liganded cysteines were identified *in vitro* compared to *in situ* drug treatment (298 vs. 134), 82% of *in situ*-identified sites (110 out of 134) were identified in both conditions (Figure 3b). Based on these findings, we compiled cysteine targets of FDA-approved TCIs^33^, off-targets of lead-like/FDA-approved inhibitors^18,34–37^ and immunomodulatory elaborated electrophiles^19^, showing significant enrichment for exposed cysteine residues compared to the distribution of cysteines detected by MS-ABPP or the proteome average (Figure 3c).

**Figure 3:**
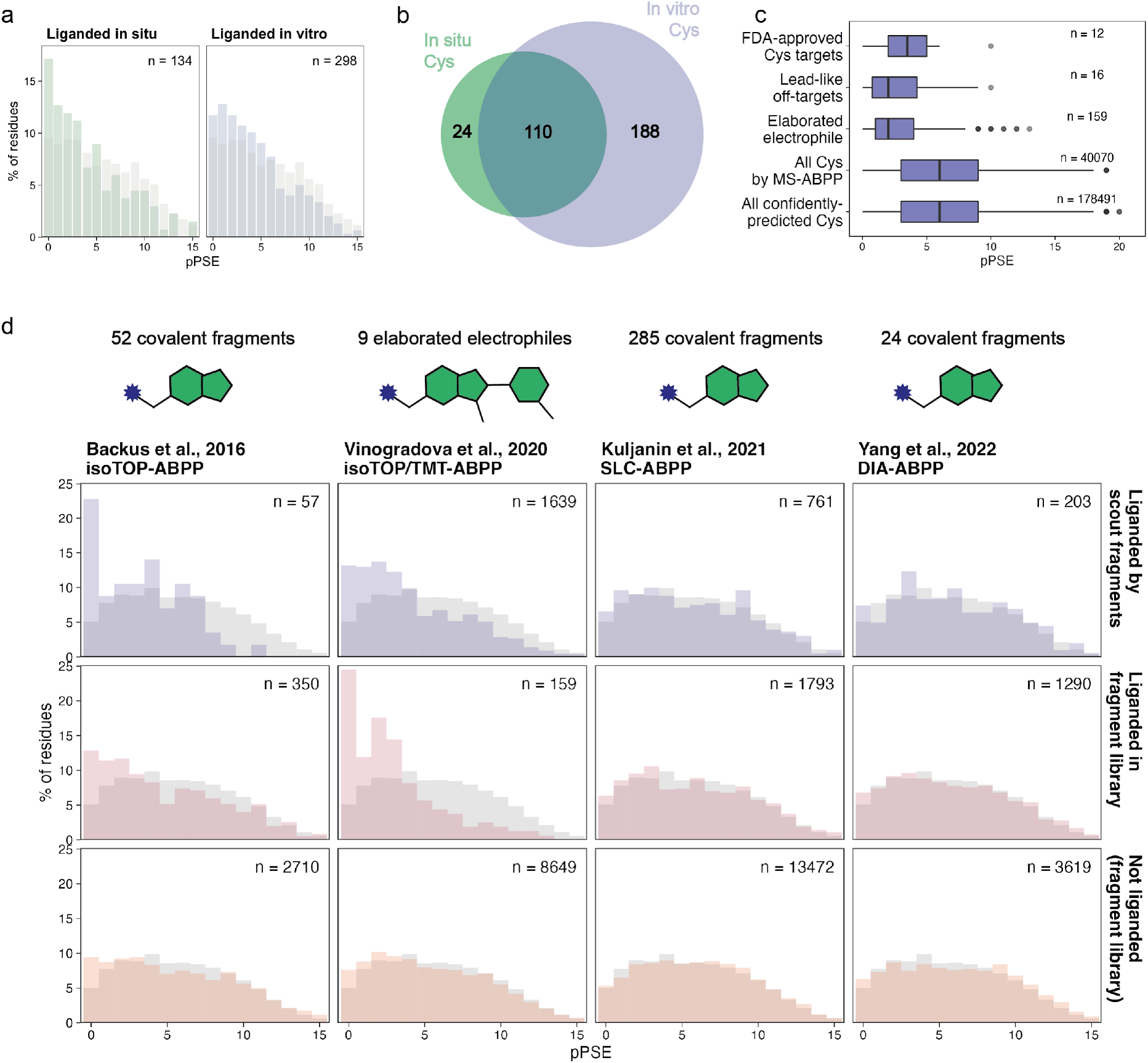
Covalent inhibitors, from small fragments to elaborated drug-like molecules, predominantly target exposed cysteines. **a.** Covalent fragment-targeted cysteines are enriched for accessible cysteines over non-liganded residues (light grey), especially when drug treatment is performed in live cells (*in situ*, left panel). Data from Backus et al.^2^ **b.** Venn diagram showing the overlap between cysteines targeted (R_DMso/Fragment_ > 4) by *in situ* and *in vitro* fragment treatments. **c.** Cysteine targets of FDA-approved TCIs, off-targets for lead-like compounds and immunomodulatory elaborated electrophiles (Vinogradova et al.^19^) show significant bias towards accessible cysteine residues, compared to the distributions sampled by all described MS-ABPP datasets and the average of all cysteine residues across the proteome. Boxplots display 25th, 50th (median) and 75th percentile, whiskers display upper/lower limits of data. Outliers are plotted as points. **d.** Comparison of accessibility distributions for four fragment screening datasets.^2,18–20^ Panels show the subset of cysteine residues liganded by two promiscuous scout electrophiles (dark blue, top panels), covalent fragment/inhibitor libraries (red, middle panels) and those not liganded (orange, bottom panels). Inset shows number of unique cysteine residues plotted in each histogram.

We finally sought to understand the structural context of cysteines liganded in large-scale competition fragment experiments. A specific set of small, promiscuous electrophilic compounds, also termed ‘scout fragments’ by analogy to the low molecular weight fragments applied in conventional fragment-based ligand discovery, have been applied in diverse biological contexts to determine differential proteome reactivity, for example upon activation of T-cells^19^ or NRF2-knockdown in non-small cell lung cancer cells.^38^ We therefore integrated reactivity data from two commonly profiled promiscuous fragments (KB02, KB05, Figure S1a) across 4 generations of CP technologies.^2,18–20^ Two datasets (Backus et al. and Vinogradova et al.) show that exposed residues are significantly enriched in scout-reactive cysteines but not non-liganded cysteines (Figure 3d, left columns). This enrichment is observed to a significantly lesser extent in the Kuljanin et al. and Yang et al. datasets (Figure 3d, right columns), perhaps in part due to predominantly non-overlapping cysteines sampled in each approach (Figure S1b,c,d). A similar trend is also evident in respective larger fragment screening experiments, where Backus et al. and Vinogradova et al. show enrichment for more exposed residues, however such enrichment is less apparent in the Kuljanin et al. and Yang et al. datasets. Categorising the liganded cysteines by warhead (chloroacetamide vs. acrylamide) from each fragment library showed no clear accessibility trends by warheads across the three datasets (Figure S2a). Furthermore, we calculated various physicochemical properties of the screened fragments (cLogP, cLogS, H-bond donors, H-bond acceptors, molecular weight, absolute and relative polar surface area, total surface area).^39^ Interestingly, highly promiscuous fragments (>50 liganded residues) predominantly came from the upper quartile of cLogP values (and lower quartile of cLogS values), perhaps indicating promiscuity at least partially results from nonspecific interactions of lipophilic fragments with protein surfaces (Figure S2b). Taken altogether, our results suggest a subset of exposed cysteines overwhelmingly represent the interacting residues of covalent fragments and, to a greater extent, more developed TCIs.

### Implications for cysteine profiling in targeted covalent inhibitor development

For applications such as target identification from phenotypic covalent fragment screening or off-target profiling of developed TCIs, comprehensive (>50%) coverage of ligandable cysteines will be required to achieve a reasonable success rate using CP workflows alone. Given inherent limitations in a bottom-up proteomics strategy (constraints on unique/detectable peptide length using only tryptic digestion, variable ionisation efficiency, proximal PTMs on a given cysteine-containing peptide), we considered the theoretically achievable coverage of druggable cysteines in a MS-ABPP experiment as a function of number of cysteines and selectivity for druggable residues. We defined selectivity as the fraction of cysteines sampled in a given experiment that are druggable compared to the proteome-wide fraction of druggable cysteines. We then calculated the percentage coverage of ‘druggable’ cysteines for combinations of (theoretical) experiments quantifying up to 100,000 cysteines and achieving up to 10-fold selectivity of druggable residues. For an experiment with a selectivity value of 1, druggable cysteines are sampled in the proportion they exist in the whole proteome, therefore the percent coverage of druggable cysteines is the same as the percent of all cysteines detected. In this case, only 10% coverage is therefore achieved by profiling ~20,000 cysteines (Figure 4, dotted line). Conversely, with a 5-fold increase in selectivity at the same number of cysteines profiled, 50% of druggable cysteines would be profiled. Consistent with recent advances in high-throughput proteome profiling and instrumental advances,^40–43^ peptide detection improvements in the coming years will no doubt improve the number of cysteines profiled in a given MS-ABPP experiment. Combined with technical advances in sample preparation, acquisition and analysis to alleviate known limitations of bottom-up approaches (e.g. multiple protease strategies, PTM-aware database searching), we can therefore expect improvements in reproducible cysteine depth, however our results highlight the need to also optimise CP toward higher selectivity for potentially druggable residues.

**Figure 4:**
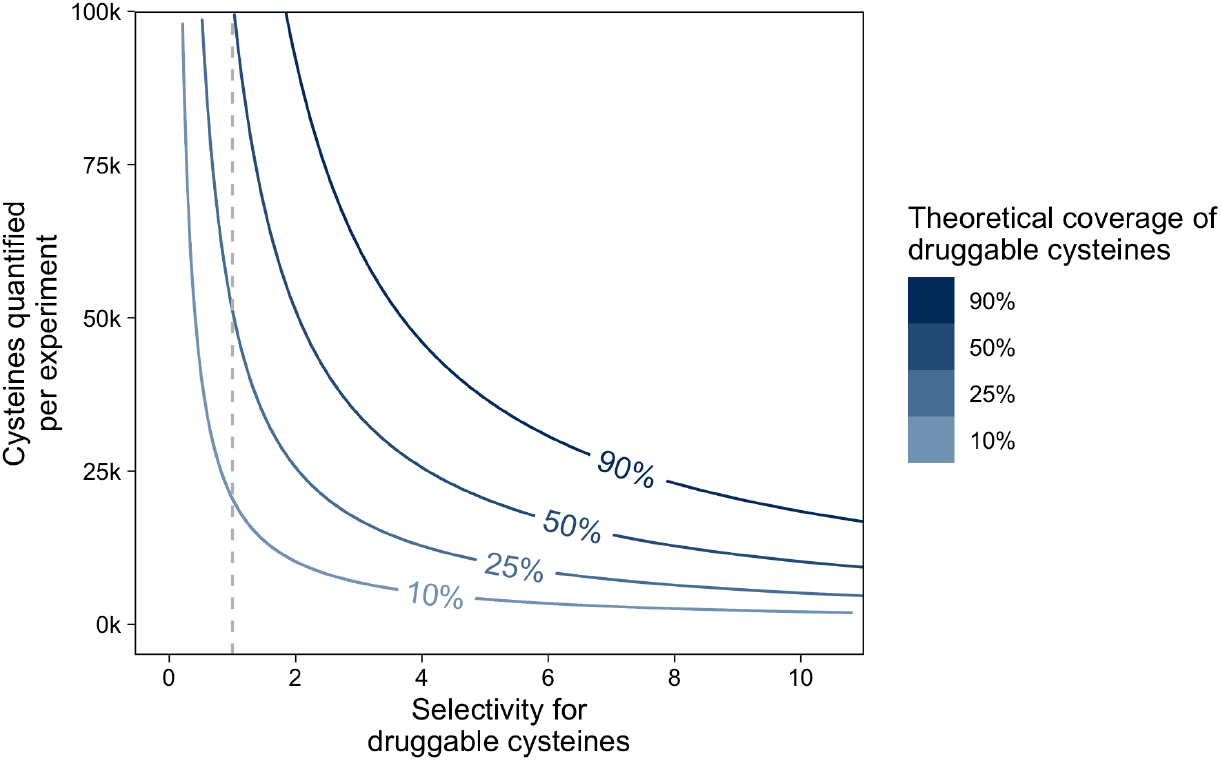
Theoretical coverage of druggable cysteine residues by MS-ABPP as a function of quantified cysteine residues per experiment (y-axis) and selectivity for druggable cysteine residues (x-axis). Dotted line shows the selectivity value of 1, where no enrichment of druggable residues is present.

## Conclusion

Activity-based protein profiling technologies have made an important contribution to the covalent drug discovery pipeline, with cysteine-focused competitive profiling in particular offering a method for proteome-wide dissection of cysteine reactivity and ligandability. Our meta-analysis of publicly available cysteine profiling datasets and proteome-wide accessibility predictions shows that while different warhead chemistries sample distinct accessibility distributions, there is marked consistency across datasets with similar warheads. Furthermore, photocaged α-haloketones show remarkably good performance in enriching accessible cysteines, potentially due to a combination of both a native labelling environment and dynamics of labelling afforded by photo-uncaging in live cells. We also observe significant enrichment for accessible residues in liganded targets of promiscuous scout fragments, fragment libraries and drug-like elaborated electrophiles. The early stages of covalent ligand optimisation campaigns are typically undertaken without the benefit of structure-guided design,^19,34,36^ and we suggest that proteome-wide structural accessibility analysis may be a useful complementary approach when paired with CP workflows at each step of ligand optimisation to probe the bias between accessible structural environments during evolution of a given ligand series.

Taken together, our analyses encourage further development of covalent ligand discovery workflows to enhance and optimise accessible residue coverage. Current high-throughput CP workflows spread coverage broadly across the full distribution of cysteine accessibility and appear to greatly under-sample potential ligandable cysteines, even considering recent improvements in mass spectrometry which put 25,000+ quantified cysteines within reach. However, as noted above, current MS technology is already capable of approaching full coverage of accessible cysteines if profiling capacity is tightly focused on accessible residues, for example by further development of in-cell labelling workflows driven by photocaged warheads. This consideration is particularly important for *de novo* target deconvolution of bioactive compounds with a covalent mode of action, where incomplete coverage of ligandable cysteines greatly reduces the likelihood of positive target identification. Furthermore, as binding is inferred by loss-of-signal, there is potential for false positives through compound-induced changes in proximal PTMs, redox state or protein conformation (and therefore cysteine accessibility) or simply sample handling. In these cases, direct labelling, for example with a clickable tag, is likely to be the more successful approach, through positive target enrichment.

We also identify several limitations of accessibility analysis which might be usefully addressed in the future. These include the potential for false positive or negative identification of exposed residues due to current limitations in structure prediction, which we have sought to minimise by limiting our global analysis to residues with high-confidence prediction, and limiting ligandability analysis to structured regions. Whilst we believe our approach to be as useful and robust as the underlying AlphaFold data at a whole proteome scale, any single prediction in isolation must be coupled to experimental validation to be considered actionable. The value of this analytical approach is expected to continually improve with evolution of machine learning approaches to structure prediction which take account of confounding factors, such as the presence of protein complex interfaces or PTMs which might dramatically alter accessibility. Furthermore, it is important to recognise that accessibility is not synonymous with reactivity, and it is likely that a certain percentage of accessible residues are not amenable to liganding with warhead chemistries currently employed in profiling workflows; analyses will therefore benefit from refinement in parallel with the disclosure of increasingly large and diverse profiling datasets which reflect broader residue level reactivity. A tighter integration with proteomic data might be achieved in future using filters which account for the limitations of proteomic analysis, for example the limited capability of proteomics workflows to deal with very short, very long, or highly modified peptides, encompassing a significant proportion of cysteines which may be important ligand target sites.^44^

Finally, we note that the analytical pipeline presented here is straightforward to implement, and should be equally applicable to any residue-specific profiling pipeline. For example, as new profiling datasets become available it will be interesting to observe the evolution of reactive amino acid coverage across models and species enabled by the 200 million predicted protein structures in the AlphaFold2 database, and by ongoing developments in warhead chemistries targeting residues beyond cysteine.^4,45^

## Methods

### pPSE prediction from human AlphaFold2 structures

Solvent accessibility prediction, intrinsically-disordered region prediction and secondary structure annotations were calculated using StructureMap as described in Bludau et al.^21^ (https://github.com/MannLabs/structuremap). Jupyter Notebooks to reproduce the analysis are publicly available at www.github.com/TateLab.

### Data curation and filtering

All datasets were downloaded as Supplementary Information files from their respective publications (Table S1). For analysis of cysteines detected in each dataset, all unique cysteine residues were extracted and matched to AlphaFold-predicted pPSE values by UniProt accession ID and sequence position. Any reverse/contaminant proteins and proteins with no predicted pPSE values were removed and cysteines were then filtered by the following criteria: ambiguous cysteine identifications (i.e. peptides/annotations with >1 cysteine) and low-confidence predictions (Alphafold prediction quality < 70) were removed. Additionally, for fragment ligandability analysis (Figure 3), cysteines predicted to be in unstructured proteins domains were removed.

### pPSE distribution analysis

For all distributions, the number of residues at each pPSE value was normalised by the total number of cysteines per experiment/condition such that the sum of all pPSE values was 1. All confidently predicted cysteine residues (quality > 70) were used for reference distributions (light grey, Figure 2a, 2e, 3a, 3d),

### Ligandability analysis

Fragment screening datasets were downloaded as Supplementary Information files from their respective publications (Table S1). Integration with AlphaFold-predicted pPSE values and subsequent filtering was performed as described above. Only fragment-cysteine interactions with R_DMso/Fragment_ > 4 were annotated as liganded.

### Fragment property analysis

SMILES strings were manually collated from Supplementary Information files of each fragment screening dataset and imported into OSIRIS DataWarrior^39^ (v 5.5.0, Actelion Ltd). Physicochemical properties were calculated using DataWarrior built-in functions and exported to R for visualisation.

### Theoretical coverage calculations

The percentage coverage of druggable cysteines was calculated at each discrete combination of experimental coverage (N_Detected Cys_) and selectivity by the following equation:

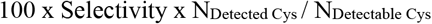

We based our calculation on a total of 204,707 detectable cysteines from an *in silico* digest of 19776 human proteins by Yan et al.^28^ The calculations were implemented in the R statistical environment.

## Data/code availability

All published datasets are available from their respective publications (Table S1). R scripts for re-formatting and collating literature data sources, analysis and generation of all figures are freely available at www.github.com/TateLab. All code along with requisite raw and processed data are also available together at https://data.mendeley.com/v1/datasets/ysvwmmmsxv/draft?preview=1

## Acknowledgements

Figure 1b/c were created with BioRender.com. We are grateful to members of the Tate group at Imperial College and the Francis Crick Institute for their advice and suggestions during the refinement of this manuscript.

## Competing interests

EWT is or has been employed as a consultant or scientific advisory board member for Myricx Pharma, Samsara Therapeutics, Roche, Novartis and Fastbase; research in his group has been funded by Pfizer Ltd, Kura Oncology, Daiichi Sankyo, Oxstem, Exscientia, Myricx Pharma, AstraZeneca, Vertex Pharmaceuticals, GSK and ADC Technologies. EWT owns equity in Myricx Pharma, Exactmer and Samsara Therapeutics, and is a named inventor on patents filed by Myricx Pharma, Exactmer, Imperial College London and the Francis Crick Institute. None of these interests conflict with the work described here. J.G. has acted as a consultant for Unity Biotechnology, Geras Bio, Myricx Pharma and Merck KGaA. Pfizer and Unity Biotechnology have funded research in J.G.’s lab (unrelated to the work discussed here). J.G. owns equity in Geras Bio. J.G. is a named inventor in MRC and Imperial College patents, both related to senolytic therapies (the patents are not related to the work discussed here).

## Funding

This work was supported by Cancer Research UK with support from the Engineering & Physical Sciences Research Council (Programme Award to EWT, DRCNPG-Nov21\100001; EPSRC CDT in Chemical Biology and CRUK Convergence Science Centre PhD studentship award to MEHW, EP/S023518/1 / CANTAC721\100021 – Cancer Research UK Imperial Centre – Non Clinical Training Award). Core support from MRC (MC_U120085810) and a grant from CRUK (C15075/A28647) funded research in J. Gil’s laboratory.

## Supplementary Information

**Table S1:** Cysteine profiling datasets analysed in this work.

| **Dataset** | **Reference** | **Labelling** | **Probe chemistry** | **Enrichment chemistry** |
| --- | --- | --- | --- | --- |
| Weerapana et al. | Nature, 2010 | Lysate | Iodoacetamide | Biotin (TEV-cleavable) |
| Abo et al. | JACS, 2015 | Live-cell | Caged α-bromoketone | Biotin (Na<sub>2</sub>S<sub>2</sub>O<sub>4</sub>-cleavable) |
| Backus et al. | Nature, 2016 | Lysate | Iodoacetamide | Biotin (TEV-cleavable) |
| Abo et al. | ChemBioChem, 2016 | Live-cell | Caged α-iodoketone | Biotin (Na<sub>2</sub>S<sub>2</sub>O<sub>4</sub>-cleavable) |
| Vinogradova et al. | Cell, 2020 | Lysate | Iodoacetamide | Biotin / desthiobiotin |
| Zanon et al. | ChemRxiv, 2021 | Lysate | Benziodoxolone | Desthiobiotin |
| Cao et al. | Anal. Chem, 2021 | Lysate | Iodoacetamide | Biotin |
| Yan et al. | ChemBioChem, 2021 | Lysate | Iodoacetamide | Biotin |
| Kuljanin et al. | Nature Biotech., 2021 | Lysate | Iodoacetamide | Desthiobiotin |
| Yang et al. | JACS, 2022 | Lysate | Iodoacetamide | Biotin (acid-cleavable) |
| Kemper et al. | Nature Methods, 2022 | Lysate | Iodoacetamide | Desthiobiotin |
| Yang et al. | Curr. Res. Chem. Bio., 2022 | Lysate | $\alpha,\beta$-unsaturated ketone | Biotin (TEV-cleavable) |
| Lai et al. | ChemRxiv, 2022 | Lysate | N-Acryloylindole | Desthiobiotin |

**Figure S1:**
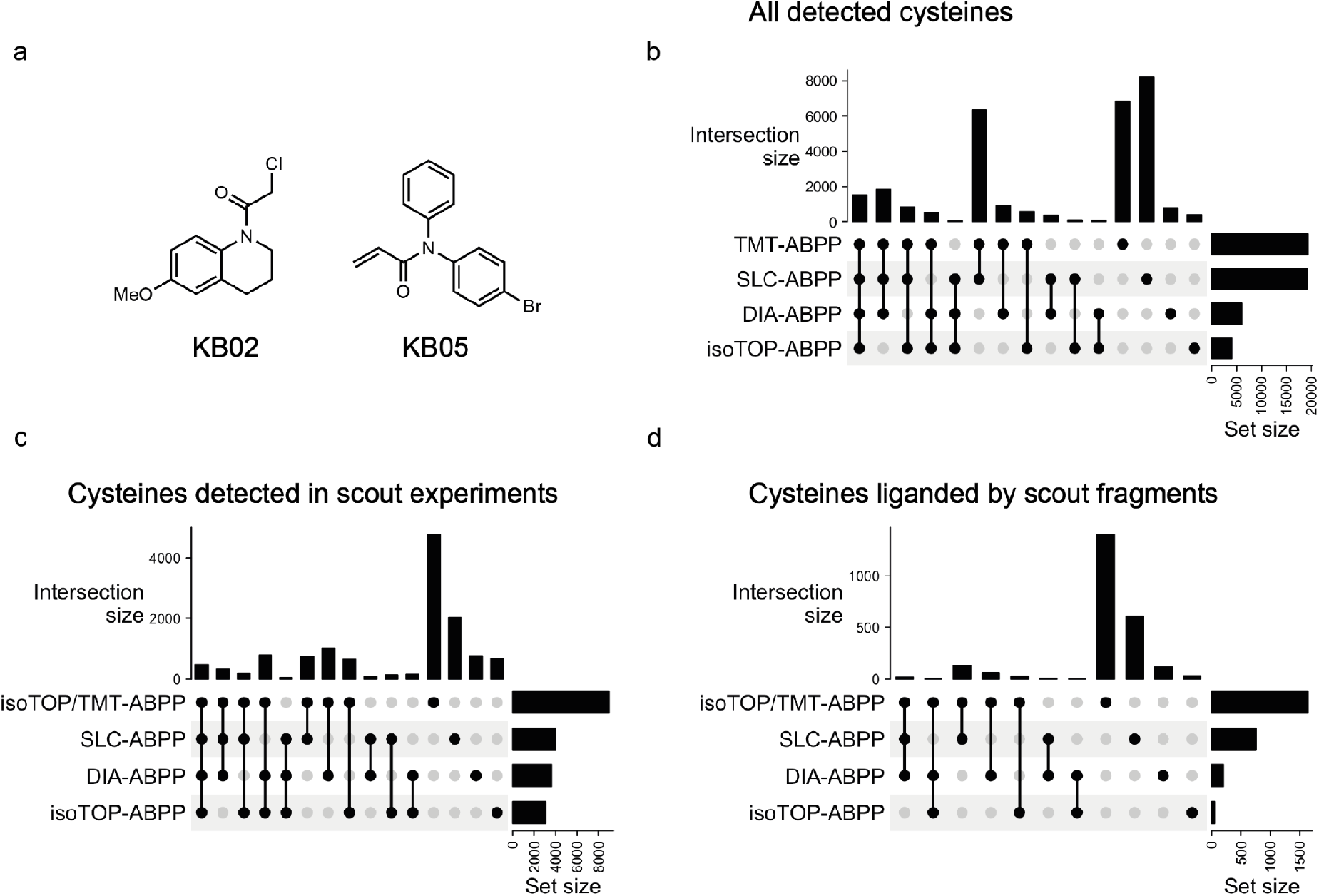
Data relating to scout fragment experiments. **a.** Chemical structures of two scout fragments (KB02, KB05) analysed across four different CP workflows. **b.** Upset plot showing overlaps of all identified unique cysteine residues across each dataset. **c.** Upset plot showing overlaps of cysteines detected in scout fragment experiments across four datasets. **d.** Upset plot showing overlaps of liganded cysteines by KB02 or KB05 in scout fragment experiments.

**Figure S2:**
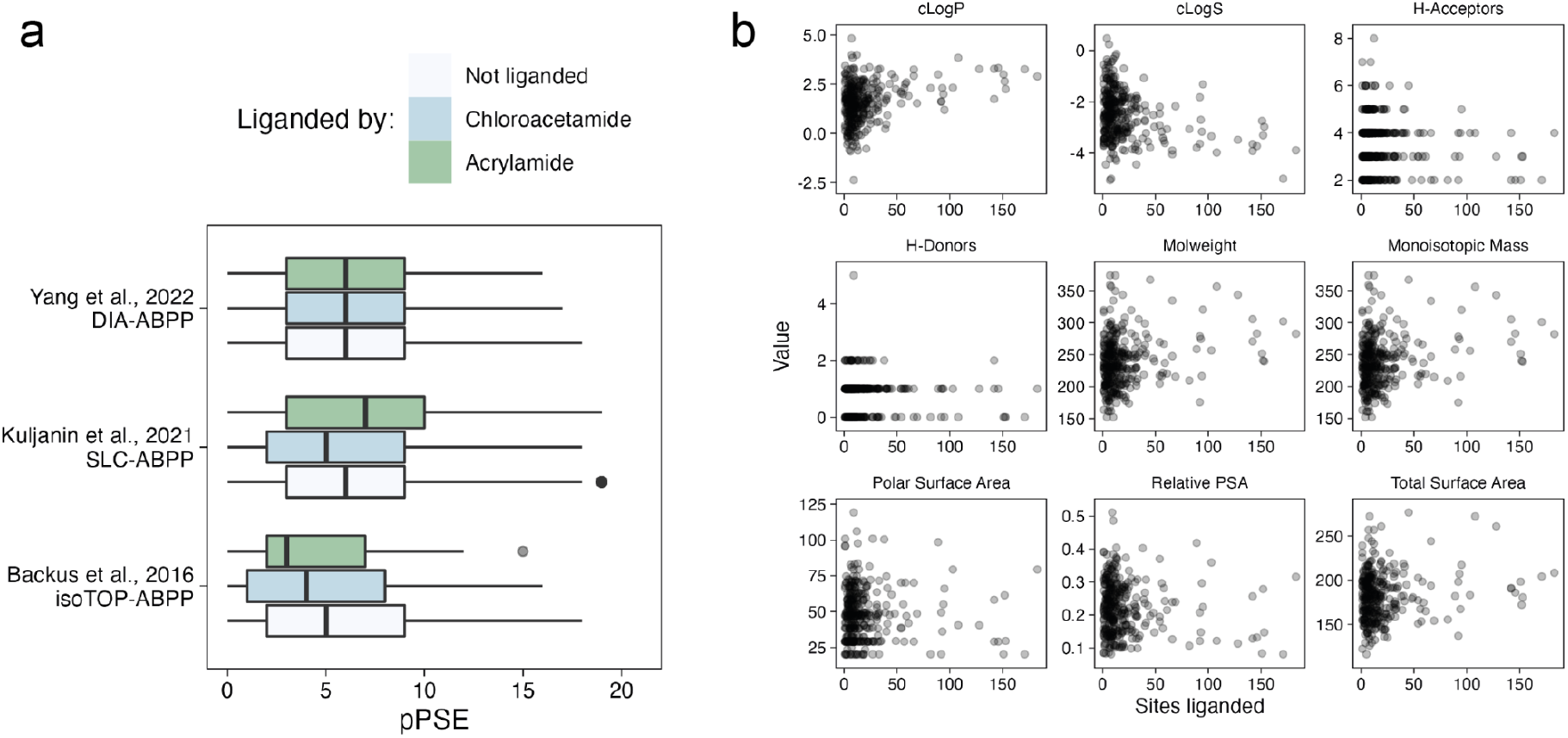
Characterising warhead-specific behaviour of fragment libraries screened by cysteine profiling. **a.** Boxplots showing distribution of pPSE for cysteines liganded either by chloroacetamide/acrylamide fragments or not liganded across three fragment screening datasets. **b.** Physicochemical properties of covalent fragments plotted against promiscuity (number of cysteines liganded by each fragment).

